# Classification of Parkinson’s Disease and Delineating Progression Markers from the Sebum Volatilome

**DOI:** 10.1101/2023.03.01.530578

**Authors:** Caitlin Walton-Doyle, Beatrice Heim, Eleanor Sinclair, Sze Hway Lim, Katherine A Hollywood, Joy Milne, Evi Holzknecht, Ambra Stefani, Birgit Högl, Klaus Seppi, Monty Silverdale, Werner Poewe, Perdita Barran, Drupad K Trivedi

**Affiliations:** Manchester Institute of Biotechnology, Department of Chemistry, University of Manchester, Manchester, M1 7DN, UK; Department of Neurology, Innsbruck Medical University, Innsbruck, Austria; Department of Neurology, Salford Royal Foundation Trust, Manchester Academic Health Science Centre, University of Manchester, Manchester, UK, M6 8HD

## Abstract

Parkinson’s Disease (PD) has been associated with a distinct odour, which emanates from the skin and is strongest in sebum-rich areas. In this study, sebum was sampled from participants using cotton gauze and the volatile components emanating from these swabs were analysed directly with thermal desorption gas chromatography - mass spectrometry (TD GC-MS). We analysed subjects with clinically established PD (*n*=46) along with healthy controls (*n*=28) sampled from two sites. The volatilome profiles obtained for PD and control cohorts were compared with the profile of participants (*n=9*) with isolated REM sleep behaviour disorder (iRBD) to investigate metabolite changes in probable prodromal PD. We also compared PD participants sampled at yearly intervals for a total of three years. Volatile compounds from TD GC-MS analysis were found in different quantities between PD, control and iRBD subjects. We found 55 significant features where abundance in samples from individuals with iRBD was intermediate between that found for PD and control samples. Significant features were found to be alkanes and fatty acid methyl esters (FAMEs), with other metabolites identified as an aldehyde, purine and tropinone. In olfactory analysis of the iRBD samples three out of nine were classified PD, and on clinical follow up two of these showed PD symptoms. Further, when analysing the volatilome from longitudinal PD sampling, almost two-thirds of the significant features showed differential regulation over the three visits. Our findings support the use of sebum as an accessible biofluid rich with measurable volatile compounds which alter in abundance in individuals with PD and iRBD, as the disease progresses.

## Introduction

Parkinson’s Disease (PD) is the second most common neurodegenerative disorder, with more than six million people affected worldwide and an expected two-fold increase in prevalence over the next generation.^1^ The pathological hallmarks are neuronal α-synuclein positive inclusions and cell loss in the substantia nigra, other areas of the brain, and the peripheral autonomic nervous system causing the cardinal motor features of bradykinesia, rigidity and rest tremor as well as a plethora of non-motor symptoms.^2^ Current diagnostic criteria for PD are anchored on clinical features, which overlap with other conditions and may result in initial misclassification.^3^ Highly sensitive and specific tests, therefore, are a major need in PD - both to enhance diagnostic accuracy and to enable detection of the earliest disease stages where future disease-modifying therapies would be the most effective.^3^

Clinically manifested PD evolves from a prodromal stage characterised by the occurrence of a variety of non-motor symptoms, including anosmia, constipation, anxiety, depression, change in cognitive behaviour as well as REM-Sleep Behaviour Disorder (RBD).^4,5^ Together with various risk markers, these have been used to define research criteria for prodromal PD. Within these criteria a polysomnographic diagnosis of isolated RBD (iRBD), characterised by loss of REM-atonia and jerks or dream-enacting behaviours during REM sleep confers the highest risk for future PD with a likelihood ratio of 130.^6^ In order to diagnose RBD, polysomnography (PSG) is needed to demonstrate the presence of REM sleep without atonia, which is an excessive electromyographic activity during REM sleep.^6^

PD has been associated with a distinct odour, strongest in sebum-rich areas. Sebum is a lipid-rich substance secreted by the sebaceous glands on the skin, which are most concentrated on the face and upper back. Increased sebum output (seborrhoea) and seborrheic dermatitis are well-known features of PD and changes in volatile profiles of sebum have recently been detected using Thermal Desorption – Gas Chromatography – Mass Spectrometry (TD GC-MS).^7,8^ Sebum is an alternative biofluid found on the skin and that can be obtained by non-invasive sampling, giving rise to advantages such as at-home sampling as well as low consumable and transport costs.. It has been shown sebum does not require the same cryogenic storage as other biofluids to maintain its diagnostic capabilities further reducing costs associated.^9^

To date, TD GC-MS has not been employed on cohorts showing prodromal PD symptoms, despite evidence that the change in odour occurs premotor symptoms. This motivates our investigation to determine if volatile components of sebum have potential for biomarkers for early diagnosis. In this study, we analysed sebum samples from PD and control cohorts finding good discrimination between their volatile metabolite profiles adding to our body of work in this area and improving our discrimination between diseased and control. We investigated features that contributed highly to the separation of phenotypes and putatively identified them using fragment spectra. We further analysed samples from iRBD participants. We describe how odour and mass spectrometry markers permit classification between these three different phenotypes, and features where iRBD concentration lies between disease and control, indicative of possible biomarkers of early stages of disease. Further, we examined markers in samples collected longitudinally from PD participants at three annual timepoints to investigate possible progressive features.

## Subjects and Methods

### Sample Collection and Clinical Cohort

Subjects were recruited at two sites: the Movement Disorder and Sleep Disorder out-patient clinic, at the Department of Neurology, Medical University Innsbruck, Austria (Austria cohort) and the Department of Neurology at Salford Royal Foundation Trust (UK cohort). Clinic staff, patients’ caregivers, and patients without neurodegenerative or ongoing dermatological disorders were invited to participate as healthy controls. Sebum samples were collected from the upper back of participants using sterile cotton-based gauze swabs (7.5 × 7.5 cm) by single individuals to avoid variance due to sampling. The sampling protocol is described in earlier work, although in this study, samples were taken at two sites from well-studied cohorts with refined instructions and identical for sampling..^7,8^

The UK cohort consisted of people with PD (PwPD) (*n*=30) and control (*n*=19) participants collected under NHS Research Authority REC reference: 15/SW/0354. From 30 PD participants, 11 were sampled longitudinally three times in total, ∼12 and 24 months after the first sampling. Controls were not included in longitudinal sampling. The Austrian cohort consisted of participants diagnosed as PwPD (*n*=16) and PSG-confirmed iRBD (*n*=9)^6^ as well as controls (*n*=9) collected under ethical approval from the Medical University of Innsbruck reference: AN1979 336/4.19 401/5.10 (4464a).

Participant demographics are detailed in Table 1, all PD participants were diagnosed by a Movement Disorder specialist, using the UK Brain Bank Criteria.^10^

**Table 1:** Participant demographic information and sample groups for analyses. For age and BMI, standard deviation (SD) is shown in parentheses.

| **Location/phenotype** | **Participants** | **Sex**<br>**(M:F)** | **Age**<br>**mean (SD)** | **BMI**<br>**mean (SD)** |
| --- | --- | --- | --- | --- |
| UK/ PD Timepoint 1 | 30 | 5:1 | 63.83 (7.87) | 27.01 (3.67) |
| UK/ PD Timepoint 2 & 3* | 11 | 11:0 | 60.09 (8.51) | 27.42 (2.78) |
| UK/ Controls | 19 | 10:9 | 63.89 (11.19) | 26.37 (3.87) |
| Austria/ PD | 16 | 3:1 | 64.63 (6.38) | 25.77 (3.78) |
| Austria/ iRBD | 9 | 8:1 | 69.89 (11.83) | 27.93 (4.69) |
| Austria/ Controls | 9 | 8:1 | 60.44 (4.80) | 27.06 (3.61) |

\*consisted of same 11 PD participants sampled at ~12 and 24 months after first sampling

**Table 2:** Description of three models generated to compare phenotypes and samples included in each. For longitudinal analysis the age stated is based on that at the first visit. For age and BMI standard deviation (SD) is shown in parenthesis. For gender, male-to-female ratio is shown in parenthesis.

| **Comparison** | **PD** | **iRBD** | **Control** |
| --- | --- | --- | --- |
| *PwPD vs. Control*<br>(Fig 1) | Age 64.11 (7.32)<br>BMI 26.58 (3.72)<br>Gender (37:9) | N/A | Age 62.79 (9.64)<br>BMI 26.58 (3.72)<br>Gender (18:10) |
| *PwPD vs. Control vs. iRBD*<br>(Fig 2) | Age 64.63 (6.38)<br>BMI 25.77 (3.78)<br>Gender (3:1) | Age 69.89 (11.83)<br>BMI 27.93 (4.69)<br>Gender (8:1) | Age 60.44 (4.8)<br>BMI 27.06 (3.61)<br>Gender (8:1) |
| *PwPD progression over 24 months since first sample* (Fig 3) | Age 60.09 (8.51)<br>BMI 27.42 (2.78)<br>Gender (11:0) | N/A | N/A |

Two iRBD subjects developed subtle parkinsonian signs after follow-up but had been free of any such sign at the time of sampling. The majority of our PD and iRBD participants are male in line with the global prevalence rates.^11,12^

All participants provided informed consent for this study.

Following collection, the gauze swabs were placed in Ziplok plastic bags, transported to the test site and stored at −80 °C until analysis. It has previously been shown storing samples at ambient temperatures does not affect the capability of classifying PD, and thus the time spent out of cryogenic storage in collection and postage is not considered damaging to the validity of the samples here.^9^ Dynamic headspace was analysed from all these samples in a randomised order. Raw data were deconvolved, annotated and converted to a data matrix consisting of 613 features. We interrogated these data. In summary, we sought to answer two questions:

1. Do components that contribute to the skin swab VOC signature alter as Parkinson’s progresses?
2. Is there an VOC signature associated with iRDB and can it be distinguished from that of PD and of control participants with TD GC-MS?

## Materials

Sterile gauze swabs (Arco, UK) and sample bags (GE Healthcare Whatman, UK) were used for all sample collection. The headspace was collected into 20mL headspace vials (GERSTAL, Germany) and absorbed on to TENAX TA sorbent tubes and packed liners (GERSTAL, Germany). The solvents used for making system suitability mix were Optima LC-MS grade methanol (Fisher Scientific) and CHROMANORM absolute 99.8% ethanol (VWR Chemicals). The system suitability mix used to check for instrumental drift was a mix of seven compounds (Sigma-Aldrich): l (-)-carvone (27.0 μM), δ-decalactone (96.4 μM), ethyl butyrate (150.5 μM), ethyl hexanoate (30.5 μM), hexadecane (43.7 μM), nonane (83.07 μM), and vanillin (100.8 μM), all in MeOH: EtOH (9:1).

### Analytical Setup

Samples were thawed and removed from bags before transference to headspace vials (20mL). Analysis was performed on an 7890A GC (Agilent) coupled with a 5975 MSD (Agilent) equipped with a GERSTAL MPS handling system. Volatile analytes were preconcentrated by incubating samples at 80°C for 10 mins. Analytes were trapped using 1L dry nitrogen purged through the headspace at 70 mL/min and onto the TENAX TA sorbent tube held above at 40°C. The sorbent tube subsequently was transferred to the Thermal Desorption Unit (TDU) located in the GC inlet where it was held at 30°C for 1 min before the temperature increased at 12°C/s until 280°C and held for 5 min. The Cooled Injection System (CIS) was held at 10°C for 0.5 min before a 12°C/s ramp up to 280°C and held for a further 5 min. Analytes were separated on an Agilent VF-5MS column (30m x 0.25mm x 0.25um) with a flow of 1mL/min and a split ratio of 1:20. The temperature of the oven was held at 40°C for 1 min then ramped at 25°C/min to 180°C, 8°C/min to 240°C and held for 1 min then 20°C/min to 300°C where it was held for 2.9 min. Positive pressure was added to the splitter plate at 1.6mL/min. The MSD operated in scan mode between *m/z* 30-800 with the source temperature held at 230°C and quadrupole temperature held at 150°C.

### Olfactory Analysis

Samples (gauze swabs in plastic bags) of known phenotype were presented to the tester (an individual with hyperosmia - a heightened sense of smell) to train and familiarise. Once training was completed, blinded samples were presented to the tester in the presence of observers. The tester was asked to describe the strength and type of smell, which was recorded on a ranking scale with comments. The tester ‘reset’ their nose by smelling a scent refresher of their choice between samples and periodic breaks were taken if the tester was overwhelmed. A subset of samples were revisited twice at the testers request.

### Data Processing and Statistical Analyses

Raw mass spectrometry data files were converted to .mzXML open format using ProteoWizard MSconvert.^13^ Subsequently, .mzXML files were deconvolved in R using the eRah package^14^ resulting in a data matrix with 613 deconvolved and putatively identified features. The features were assigned putative identifications using the GOLM and NIST 14.0 databases within eRah. Assignments of TMS derivatives, siloxanes or features with a match factor <70 were not considered and removed from further analyses. The final data output was a matrix of features and the relative intensity of these in each sample. In the absence of biological pooled QC, a mixture of standards was injected five times at the start of each batch, after every 5 sample injections and three times at the end of each batch. The instrument’s stability was assessed using the relative standard deviation (%RSD) of the peaks from this system suitability mix. All peaks of the SST showed %RSD <20% and data quality were considered satisfactory. All data was normalised to the sum of the total ion count, log transformed and auto scaled prior to statistical analyses. We employed multivariate statistical analyses methods to analyse the mass spectrometry data as described in supplementary information S1. Sparse partial least squares discriminant analysis (sPLS-DA) was used to model data classifying between the studied cohorts described in the table below.

### Confounders

We investigated the confounding effects for participants’ age, BMI, gender, alcohol consumption (units per week), smoking habits (cigarettes per week) and existing comorbidities. The common comorbidities within our cohorts were high cholesterol, hypertension, history of skin conditions, bone disease, thyroid conditions and heart conditions. We investigated the likelihood of any confounding effects for all these (as described in supplementary info S2).

## Results and Discussion

### PD classification using sebum VOCs (PwPD *vs*. Control)

Following deconvolution, the resultant data matrix contained 613 robust features. The sPLS-DA scores plot (Figure 1a) shows good clustering of the two phenotypes, which was cross validated and had an error rate of 0.2. We have removed the odor port from our TD-GC-MS instrument for this work, which resulted in higher signal (by one order of magnitude from our earlier work)^7,8^, and this reveals more features from the samples some of which are more robustly diagnostic. Of the 613 features, 85 were determined to be significant variables (VIP > 1, stability > 0.8) that enable classification between PD and control. When investigating regulation, 25 of the significant variables are seen to be upregulated with PD and 60 downregulated. Of the significant features, 28 could be assigned putative identifications at MSI level 2: seventeen as alkanes and seven as Fatty Acid Methyl Esters (FAMEs). Purine, tropinone, oleamide and hexadecanal were also identified and upregulated in the control samples. Lack of identification of the remaining significant features can be attributed to the lack of a suitable match in spectral libraries and absence of precursor ions, or possibly be due to chromatographic co-elution. Example expressions of annotated significant features are displayed in Figure 1b. This data validates our earlier observation that volatile compounds from sebum can distinguish PD from control, irrespective of geographical location of subjects. A list of all features found significant is shown in supplementary information table S1.

**Figure 1:**
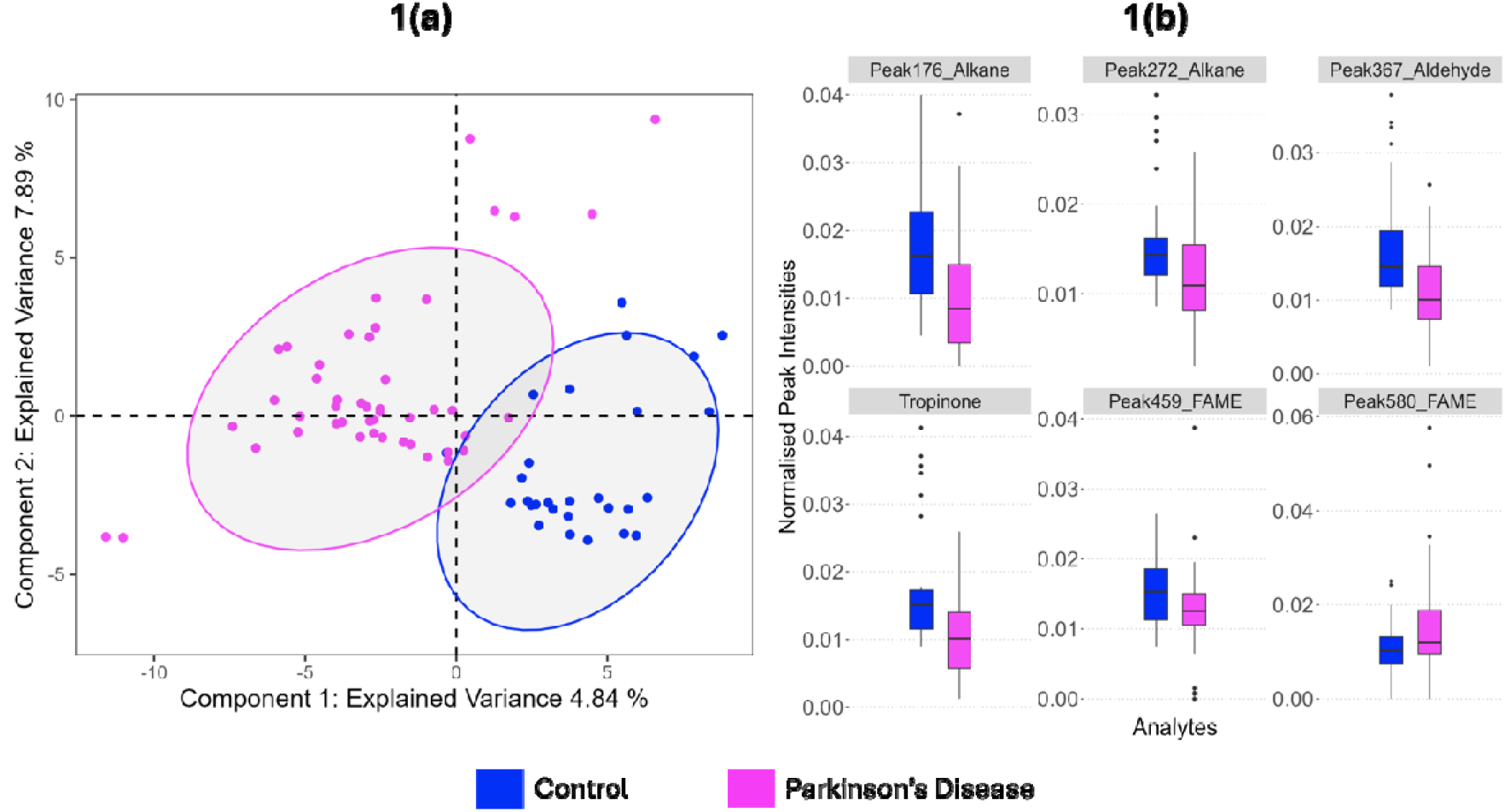
Modelling VOC profiles from skin swabs of Parkinson’s disease and control participants. (a) Optimised sPLS-DA showed two clusters distinctly separating PD and controls, validated by 3-fold cross validation with 100 repeats. (b) Example box plots for six of the 85 significant features, chosen to demonstrate changes in mean expression in PD and controls, selected using VIP scores (>1) and cross validation feature stability scores (>0.8). For further details on the selection of these features see supplementary information.

### Classification of iRBD from control and PD

We examined the cohort collected from Austria using all 613 features from the data matrix. The scores plot of sPLS-DA (Figure 2a) shows 3 distinct groups (cross validated with an error rate of 0.06). The control participants cluster the most tightly and thus show the least variability between individuals, while more within-group variability is seen in iRBD and PD. Examples of putatively identified features displaying mean iRBD regulation between that of PD and control groups are displayed in Figure 2b.

**Figure 2:**
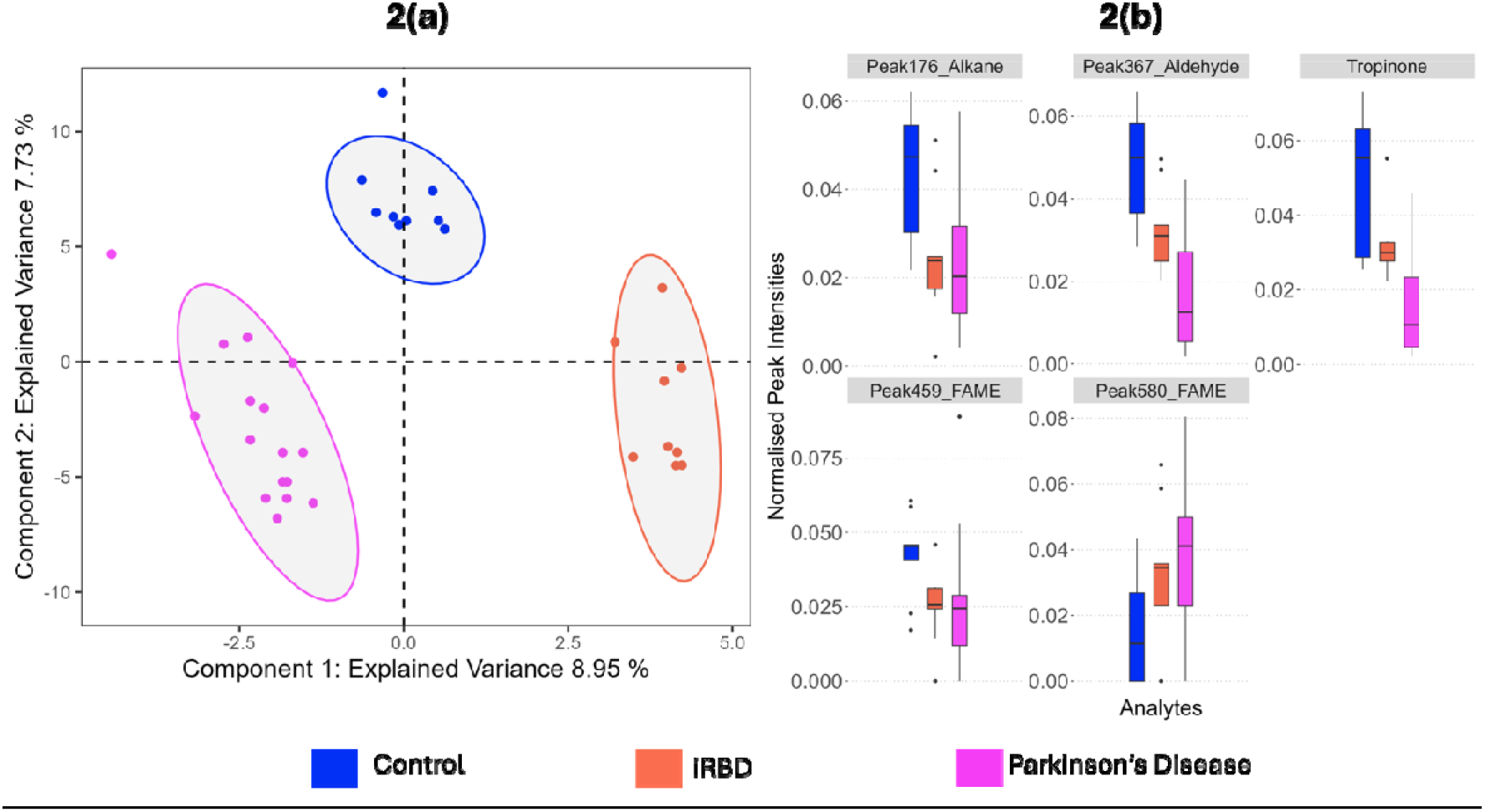
Modelling VOC profiles collected from skin swabs of PD, control and iRBD participants from the Austrian cohort. (a) Optimised sPLS-DA showed three clusters distinctly separating PD, controls and iRBD participants validated by 3-fold cross validation with 100 repeats. (b) Examples of significant features selected using VIP scores (>1) and cross validation feature stability scores (>0.8) to compare mean expression in PD, controls and iRBD, are shown with box plots. For further details on the selection of these features see supplementary information.

Of the 613 features, 76 features had VIP score > 1 (within component 1 and 2) and cross validation feature stability score > 0.8 and thus were considered significant. Out of these 76 features, 55 showed the mean iRBD concentration to be between that of PD and control, suggesting possible progression features. Nineteen of these 55 possible progression features could be putatively identified (MSI level 2) and these were: 7 alkanes (different regulations), 8 FAMEs (different regulations), two annotations of tropinone (upregulated in control), an aldehyde (upregulated in control) and purine (upregulated in PD). Five of those annotated features are shown in Figure 2b.

### Odour Analysis of iRBD

All iRBD samples along with three PD and three control samples were subjected to odour analysis as described previously.^15^ The hyperosmic individual (HI), who was not told how many of each phenotype was presented, was able to distinguish PD from control in all cases and also classify six of the iRBD samples separately as possessing a distinct odour. Three of the iRBD samples were assigned as PD by the HI. Two out of the three participants classified as PD showed signs of conversion to PD at clinical follow ups post sampling. This observation indicates that the odour of each disease can be distinguished and that iRDB has a distinct volatilome containing volatile markers that may report on the conversion/progression to PD.

### VOC signature with Parkinson’s disease progression – longitudinal data

When comparing the volatile features found from the same individuals sampled at yearly intervals, the skin volatilome exhibits good separation determined by cross validated (error rate of 0.15) sPLS-DA (Figure 3a). The intragroup variation increases over time, with tighter clustering seen at the first year of sampling, indicating higher variation between individuals as PD progresses. Due to the separation of the timepoints on both component 1 and 2, features with VIP > 1 in either component, as well as stability score > 0.8 were investigated and considered significant. A total of 38 features were selected for further analysis and annotation.

**Figure 3:**
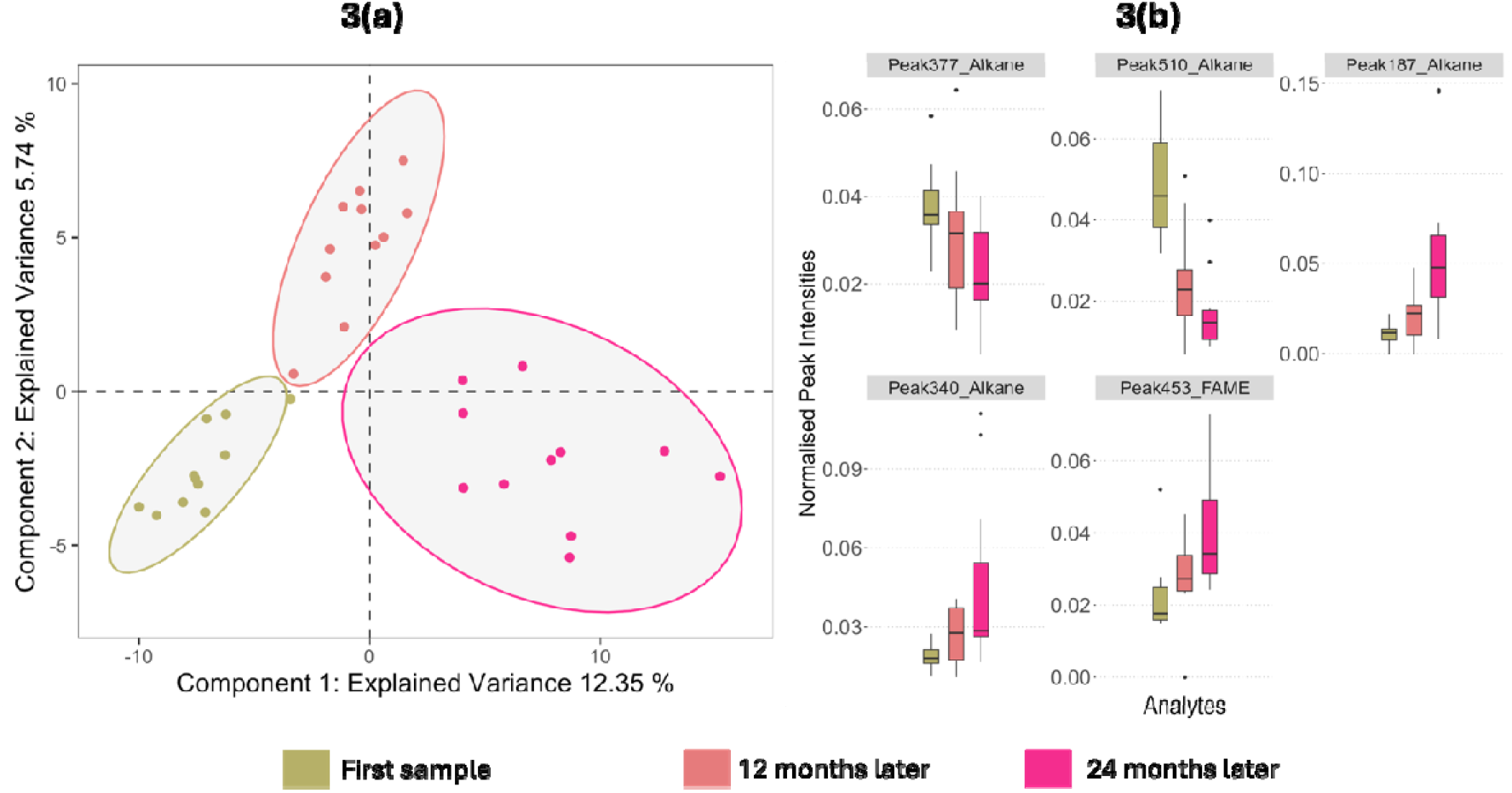
Modelling VOC profiles collected from skin swabs of PD participants from the UK cohort over three consecutive years. (a) Optimised sPLS-DA displayed three clusters distinctly separating the time points in a trajectory from left to right, validated by 3-fold cross validation with 100 repeats.(b) Example significant features that exhibited increasing or decreasing trend as time progresses; selection made using VIP scores (>1) and cross validation feature stability scores (>0.8) to compare mean expression in PD over time. For further details on the selection of these features see supplementary information.

Of the 38 significant features, 22 displayed regulation in abundance over the visits, meaning after 12 months, the abundance was between that found at the time of initial sampling and at 24 months from sampling (Figure 3b). Interestingly none of the features found as significant in classifying PD (Figure 1 and above) were significant and regulated over the timepoints, implying that the progression features are distinct from diagnostic features. Five of the features that displayed regulation could be assigned a putative identification: four as alkanes (different regulations) and one as a FAME (increasing over the three visits) (Figure 3b).

We sought to determine if these progression features were due to any confounding effects such as age, gender, BMI, medication or comorbidities. We adjusted for known confounders, including them as variables within our models. Further, we used variables to predict these confounders to ascertain the specificity of these measured metabolites to disease phenotype (Tables S1-S4). All confounders ranked below all metabolites while gender, high cholesterol and heart conditions ranked in the bottom half of the ranking list (Table S2a and S2b). Additionally, none of the models using significant features could predict confounders as an outcome (Table S3) which indicates that these significant features are specific to the phenotypes classified, not any potential confounders. We found no strong correlations (>0.5) between significant features and drug intake and the measured volatilome was not predictive of any clinical characteristic or drug intake (Table S2c and S4). These analyses overall suggest that the volatilome measured in this study is representative of the molecular phenotype of Parkinson’s disease and of iRBD and is independent of other clinical history and medication.

### Features of Interest

We investigated and interpreted the features of interest in all three models using putative annotations from spectral matches with compound libraries at metabolomics standards initiative (MSI) level 2 confidence.^16^

### Lipid Breakdown Products

Analysing lipid standards by GC-MS including headspace TD GC-MS gives rise to several resolvable features, many of which are putatively identified as alkanes.^8^ This indicates that substantial fragmentation occurs prior to GC separation.^8^ This degradation of lipid molecules may occur in either the headspace analysis (incubation and extraction) or upon introduction to the GC. The break down of lipids combined with the variation in branching and double bond position in hydrocarbon chains of lipid tail groups can result in many isomers identified as different features.^8,17^ Further, the harsh ionisation and fragmentation processes of GC-MS can result in lack of a molecular parent ion, thus molecules with hydrocarbon chains are often identified by their fragmentation fingerprint and carbon chain length can be difficult to annotate.

We therefore hypothesise that the alkanes and FAMEs found with high VIP and stability scores in models here are derived from larger lipids in sebum displaying the same degradation. Changes in fatty acid concentrations in brain tissue,^18^ blood^19^ and the gut^20^ have been associated with PD, and we have previously reported alterations in sebaceous lipids as features of interest in PD.^21,22^ Fatty acids are a well-documented product of the breakdown of larger lipid molecules such as acylglycerols.^23–25^ Similarly, aldehydes in headspace may arise due to oxidation of lipids, either endogenously or during the analysis.^26^ The inclusion of these as significant in the longitudinal study indicates lipids also alter with disease progression.

Tropinone was identified as significant in the PD *vs*. control model, and significant with regulation in the iRBD model, the latter of which had two features assigned. It was observed to decrease in PD and show iRBD as the intermediate concentration in both models. It is an alkaloid derived from putrescine.^27^ Studies in animal models have shown an important role of putrescine in skin physiology and neuroprotection.^28^ Oleamide, an endogenous fatty acid was observed to also be downregulated in PD in model 1. It is a sleep-inducing lipid known to decrease motility in animal models^29,30^ although interestingly it was not found to be a feature of interest when classifying iRBD.

Two features annotated as purine were found to be significant. One was upregulated in control in model 1, and a second feature found with intermediate expression in iRBD and downregulated in control in model 2 (Fig S1). The differences in regulation are likely due to two features with purine-like fragmentation being annotated with the same identification. We hypothesise these are purine derivatives and differences in regulation are indicative of pathway dysregulation, manifesting as an increase in some features of purine metabolism and decrease in others. One of the end products of purine metabolism is urate - an antioxidant that has been described as a risk marker for PD, and lower levels are also associated with higher severity of disease.^31–33^ Alterations in the purine pathway have been researched in an investigation for PD biomarkers, as well as in comparison with other neurological dieases.^34–36^

The overall volatilome of iRBD having many components seen as intermediary between PD and control lends further support to sebum as a reservoir for diagnostic biomarkers of PD. This is a discovery study showing great potential and will be validated with a larger sample set. As iRBD can evolve into different disorders (PD, Dementia with Lewy Bodies, Multiple System Atrophy), future work will focus on which of the iRBD-associated features observed in this study are most specific for conversion to either of these conditions.

## Conclusion and Outlook

This study demonstrates the ability of TD GC-MS to discriminate the volatilome of PD from control using sebum collected on gauze swabs followed by supervised multivariate analysis. When investigating significant features that contribute to classification, we find many metabolite features that can be attributed to lipid breakdown products, validating our previous findings.^8^

Using the same technique and analysing sebum from individuals with iRBD, we observe that a differential volatilome in these participants is present, providing potential for measurement of prodromal molecular markers. Significant features detected showed the iRBD cohort with a mean concentration between PD and control samples, indicative of disease progression. Finally, we analysed the volatile metabolites in the headspace of samples taken at yearly intervals to investigate changes over disease course and can demonstrate clear classification of samples based on stage of disease. The significant features again are putatively annotated as hydrocarbons and FAMEs supporting the hypothesis of differential regulation of lipids.

Such putatively identified markers limit the opportunity for detailed pathway analyses, but certainly furthers our hypothesis that the abundance of lipids and other metabolites captured in skin swabs from areas rich in sebum change significantly in the course of Parkinson’s disease.^8^ We have again shown that dysregulation of lipids can be observed in swabs from iRBD and in longitudinally sampled pWP. We have previously demonstrated that liquid chromatography-mass spectrometry (LC-MS) can be used to further investigate these larger molecules ^22^ and here show the power of direct analysis of the volatilome. With complementary analytical platforms we can better understand the mechanisms and pathways causing these changes as well as identity features that differentiate between disease and control which would be suitable for targeted MS analysis akin to that used routinely for newborn screening where more than 30M tests are conducted globally p/a^37^ as well as common and increasing use on mass spectrometry in drug discovery, clinical trials, microbiology, endocrinology and therapeutic drug monitoring.^38^

We have established a method using facile and non-invasive sebum sampling to demonstrate differential volatile signatures for PD, iRBD and control participants. We also describe changes in the volatilome as the disease progresses and offer a route to discover markers that could be used to inform the development and efficacy of new treatments. Together, these findings offer a novel approach for early PD classification, where current robust clinical tests are lacking as well as offering discrimination over disease progression.

## Acknowledgements

We thank Michael J Fox Foundation (grant ref:12921) and Parkinson’s UK (grant ref: K-1504) for funding this study and Community of Analytical and Measurement Sciences (CAMS) for supporting DT’s research (CAMS2020/ALect/10). This work was supported by the BBSRC (award BB/L015048/1) for instrumentation used in this work and by a DTP to CW-D (project ref. 2113640). We also thank our recruitment centres for their enthusiasm and rigor during the recruitment process. We are grateful to all the participants who took part in this study as well as PIs and nurses at the recruiting centres. Finally, we thank Waters for their continued support to the Michael Barber Centre for Collaborative Mass Spectrometry as well as the brilliant staff of the Mass Spectrometry and Separations Facility of the Faculty of Science and Engineering.

## Ethics

Ethical approval for this project (IRAS project ID 191917) was from the NHS Research Authority (REC references: 15/SW/0354 and from the Medical University of Innsbruck reference: AN1979 336/4.19 401/5.10 (4464a). Informed consent was received from all participants prior to their enrolment in the study.

## Data Availability

Raw data sets generated during the current study are available from MetaboLights Repository https://www.ebi.ac.uk/metabolights/search with the study identifier MTBLS11712.

