## Supplementary Information for "Classification of Parkinson’s Disease and Delineating Progression Markers from the Sebum Volatilome"

S1: Statistical analyses method description

In pilot metabolomics studies the sample size is often smaller than the number of features detected by high resolution methods used and the data suffers from high degree of multicollinearity. Instead of feature selection prior to classification task, sPLS-DA allows us to perform classification and variable selection in a single step. Clustering was determined based on scores plot and the performance of the model then was determined by comparing balanced error rates of 3-fold cross validation repeated 100 times. The number of components were selected from the initial model and then number of features per component were determined using a grid search. The final sPLS-DA model used these optimised, reduced components and features per component and the performance was then determined by balanced error rates from cross validation. Variables contributing to the classification of classes, were selected using variable importance in projections (VIP) scores and stability of the variable during cross-validation. A variable was determined to be important if VIP>1 and if stability was >0.8. These features were called *significant features* in this work. A subset that could be annotated from this feature selection was visualised using boxplots for each model.

S2: Investigation of confounding effects

To investigate if our significant features were highly ranked due to confounding effects, we performed random forest analyses with and without confounder as an independent variable. At each node attributes equal to the square root of the total number of attributes in each data were randomly drawn. We created a separate model adjusting for one confounder at a time and compared it with original unadjusted model. We repeatedly (100 times) split data in each model where 66% observations were used for training and 34% were used for testing. The final output is based on the majority vote from individually developed trees over 100 repeats on test sets. When a model’s classification accuracy improved by 10% or more in the adjusted model, a confounding effect was concluded. To account for chances of a smaller sample size or class imbalance within these modes, we limited the depth of individual trees to three, no subset splits smaller than 5 were allowed for any model and we attributed class weights inversely proportional to their frequencies. We also used each confounder as a dependent variable to determine if these significant variables can classify or regress against the confounder predicting it as an outcome. We used Matthew correlation coefficients (MCC) and regression coefficients (R^2^) to determine if the significant features in all the above comparisons, were confounded. The MCC provides a balance measure accounting for true and false positives as well as negatives from the model, even if the classes are imbalanced. The value of +1 indicates a perfect prediction, 0 indicates prediction as good as a random and -1 indicates complete disagreement between observed and predicted class. We saw no significant improvement to the MCC when adjusting for confounders when classifying between any of the phenotypes as well as one year follow up (Supplementary table 2a. 2b and 2c). Moreover, when each features’ contribution was ranked from random forest classification, none of the tested confounders were ranked within the top 50% of features indicating they do not contribute to classification and have no confounding effect on the measured volatilome. We used regression coefficient (R^2^) as regression metric and Matthew Correlation Coefficient (MCC) as a metric to summarise classification accuracy of our models. For participants with Parkinson’s in the UK cohort we used random forests to determine if the measured volatilome can also determine clinical variables associated with PD (*viz.* HY stage, MOCA score, MDS_UPDRS Score, NDS, VAS, Tremor Score, PIGD Score, years since motor symptoms, years and months since clinical diagnosis and ratio between tremor and PIGD scores), medication intake or dosage of medication. We performed Pearson’s correlation analyses between medication intake and above-described clinical features and significant variables to determine if there were any associations. For participants with Parkinson’s in our Austria cohort, we used medication intake data to determine if the measured volatilome can classify PD participants by their drug intake.

**Figure S1:** Purine was found to be upregulated in control *vs*. PD, with intermediate expression in iRBD, however was not one of the most significant markers due to low stability score (<0.8) during cross validation of models (n=100).

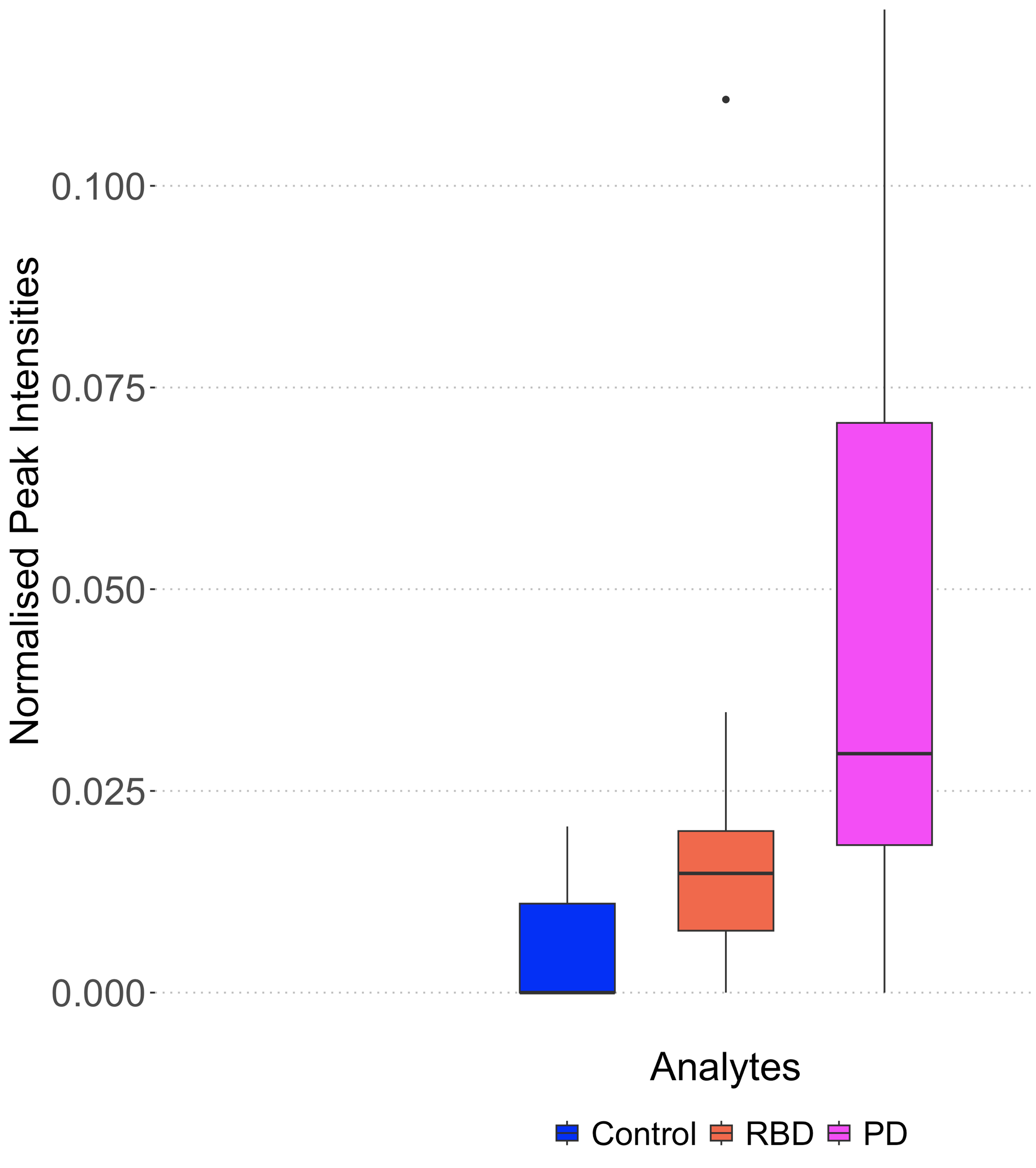

**Table S1: A list of features and annotations found significant in the three models**

| Feature | Putative ID | Found Significant in | | |
| --- | --- | --- | --- | --- |
|  |  | Model  PD v Control | Model  PD v RBD v Control | Model PD over 3 years |
| 5 | Oleamide | Y |  |  |
| 17 |  | Y | Y |  |
| 18 |  | Y |  |  |
| 21 |  | Y |  |  |
| 28 |  |  |  | Y |
| 29 | Alkane | Y |  |  |
| 32 |  |  |  | Y |
| 33 |  |  |  | Y |
| 39 |  |  |  | Y |
| 41 |  |  |  | Y |
| 46 |  |  |  | Y |
| 48 |  | Y |  |  |
| 51 |  | Y |  |  |
| 55 |  | Y | Y |  |
| 57 | Alkane | Y |  |  |
| 60 |  |  | Y |  |
| 61 |  |  | Y |  |
| 65 |  |  | Y |  |
| 66 |  |  | Y |  |
| 72 |  |  | Y | Y |
| 74 |  |  | Y |  |
| 75 |  | Y |  |  |
| 76 |  | Y | Y |  |
| 78 |  |  | Y |  |
| 79 |  |  | Y |  |
| 83 |  |  |  | Y |
| 89 |  | Y | Y |  |
| 90 |  |  | Y |  |
| 92 |  | Y | Y |  |
| 93 |  |  | Y |  |
| 94 |  |  | Y |  |
| 98 |  |  |  | Y |
| 102 |  | Y |  |  |
| 135 |  |  | Y |  |
| 147 |  | Y |  |  |
| 149 |  | Y |  |  |
| 150 |  | Y |  |  |
| 151 |  | Y |  |  |
| 152 |  | Y |  |  |
| 158 |  | Y | Y |  |
| 160 |  | Y |  |  |
| 176 | Alkane | Y | Y |  |
| 178 | Alkane | Y |  | Y |
| 179 |  | Y |  |  |
| 181 |  | Y |  |  |
| 186 | Alkane | Y |  |  |
| 187 | Alkane |  |  | Y |
| 190 | Alkane | Y |  |  |
| 193 | Alkane | Y |  |  |
| 195 |  | Y | Y |  |
| 197 |  | Y |  |  |
| 198 |  | Y | Y |  |
| 202 | Alkane | Y |  |  |
| 205 | Alkane | Y |  |  |
| 208 |  | Y |  |  |
| 209 |  |  | Y |  |
| 210 |  |  | Y |  |
| 215 |  | Y | Y |  |
| 223 |  |  |  | Y |
| 225 | Tropinone |  | Y | Y |
| 240 |  |  | Y |  |
| 248 | Alkane | Y | Y |  |
| 249 |  |  | Y |  |
| 251 | FAME |  | Y |  |
| 253 | Alkane |  | Y |  |
| 255 |  |  | Y |  |
| 257 | FAME |  | Y |  |
| 271 |  | Y |  |  |
| 272 | Alkane | Y | Y |  |
| 275 |  | Y | Y | Y |
| 276 |  | Y | Y |  |
| 280 |  | Y | Y |  |
| 286 |  | Y | Y |  |
| 287 |  |  | Y |  |
| 289 | Alkane | Y | Y |  |
| 295 |  | Y | Y |  |
| 302 |  |  |  | Y |
| 306 |  |  |  | Y |
| 310 |  | Y |  |  |
| 312 |  |  | Y |  |
| 315 | Alkane |  | Y |  |
| 327 | Alkane |  |  | Y |
| 328 |  |  |  | Y |
| 331 | Alkane |  | Y | Y |
| 339 |  | Y |  |  |
| 340 | Alkane |  |  | Y |
| 341 |  |  |  | Y |
| 343 | Alkane | Y | Y |  |
| 346 |  | Y |  |  |
| 347 |  | Y |  |  |
| 348 |  | Y | Y |  |
| 351 | FAME |  | Y |  |
| 353 |  | Y | Y |  |
| 354 |  |  | Y |  |
| 355 |  |  | Y |  |
| 358 |  |  | Y |  |
| 360 | Alkane | Y |  |  |
| 365 |  |  | Y |  |
| 366 |  | Y | Y |  |
| 367 | Aldehyde | Y | Y |  |
| 368 | Tropinone | Y | Y |  |
| 372 |  | Y | Y |  |
| 373 |  |  | Y |  |
| 377 | Alkane |  |  | Y |
| 378 |  |  | Y |  |
| 382 | Alkane |  | Y |  |
| 384 |  |  | Y |  |
| 387 |  |  |  | Y |
| 389 | Alkane | Y |  |  |
| 396 |  | Y |  |  |
| 397 |  | Y |  |  |
| 398 |  | Y |  |  |
| 399 | Alkane |  | Y |  |
| 400 |  |  | Y |  |
| 401 |  |  | Y |  |
| 402 |  |  |  | Y |
| 413 |  | Y | Y |  |
| 415 | Purine |  | Y |  |
| 417 |  |  | Y |  |
| 418 |  |  | Y |  |
| 423 | Alkane |  | Y |  |
| 426 |  |  |  | Y |
| 430 |  | Y |  |  |
| 432 |  | Y |  |  |
| 433 |  | Y |  |  |
| 437 |  |  |  | Y |
| 441 |  | Y |  |  |
| 444 | Alkane | Y |  |  |
| 447 |  |  | Y |  |
| 453 | FAME |  | Y | Y |
| 457 |  |  | Y |  |
| 458 | FAME | Y | Y |  |
| 459 | FAME | Y | Y |  |
| 471 |  | Y |  |  |
| 472 |  | Y |  |  |
| 473 |  | Y |  |  |
| 474 | Purine | Y |  |  |
| 477 |  | Y |  |  |
| 496 |  |  |  | Y |
| 497 |  |  |  | Y |
| 510 | Alkane |  |  | Y |
| 512 |  |  |  | Y |
| 513 |  |  |  | Y |
| 520 | FAME | Y | Y |  |
| 530 | Alkane | Y |  |  |
| 550 |  |  |  | Y |
| 551 |  | Y | Y |  |
| 552 | FAME | Y |  |  |
| 556 | FAME | Y |  |  |
| 559 |  |  | Y |  |
| 562 |  | Y |  |  |
| 564 |  | Y |  |  |
| 571 |  | Y |  |  |
| 579 | FAME | Y |  |  |
| 580 | FAME | Y | Y |  |
| 583 |  | Y |  |  |
| 584 |  | Y |  |  |
| 589 |  |  |  | Y |
| 592 |  |  |  | Y |
| 594 |  |  |  | Y |
| 603 |  |  |  | Y |
| 606 |  |  |  | Y |

**Table S2a: *MCC and correctly classified samples for unadjusted and confounder adjusted models comparing PD and Controls***

| Model | Overall MCC | Correct PD classification (%) | Correct Control classification (%) | Confounder in top 50% variables? |
| --- | --- | --- | --- | --- |
| Original | 0.992 | 100 | 99 | n/a |
| Age adjusted | 0.998 | 100 | 99.7 | No |
| BMI adjusted | 0.998 | 100 | 99.7 | No |
| Gender adjusted | 0.999 | 100 | 99.9 | No |
| Alcohol adjusted | 0.998 | 100 | 99.7 | No |
| Smoking adjusted | 0.998 | 100 | 99.7 | No |
| High cholesterol adjusted | 0.999 | 100 | 99.9 | No |
| Hypertension adjusted | 0.999 | 100 | 99.9 | No |
| Skin conditions adjusted | 0.999 | 100 | 99.9 | No |
| Bone diseases adjusted | 0.999 | 100 | 99.9 | No |
| Thyroid conditions adjusted | 0.999 | 100 | 99.9 | No |
| Heart conditions adjusted | 0.998 | 100 | 99.7 | No |

**Table S2b: *MCC and correctly classified samples for unadjusted and confounder adjusted models comparing PD, iRBD and Controls***

| Model | Overall MCC | Correct PD classification (%) | Correct Control classification (%) | Correct RBD classification (%) | Confounder in top 50% variables? |
| --- | --- | --- | --- | --- | --- |
| Original | 0.852 | 99.2 | 82.7 | 82 | n/a |
| Age adjusted | 0.868 | 98.8 | 85.3 | 84 | No |
| BMI adjusted | 0.868 | 98.7 | 85.7 | 84 | No |
| Gender adjusted | 0.889 | 99.3 | 88.3 | 85.3 | No |
| Alcohol adjusted | 0.876 | 99.2 | 86.7 | 84 | No |
| Smoking adjusted | 0.875 | 99 | 86.3 | 84.3 | No |
| High cholesterol adjusted | 0.895 | 99.5 | 89.7 | 85 | No |
| Hypertension adjusted | 0.891 | 99.5 | 89 | 84.7 | No |
| Skin conditions adjusted | 0.896 | 99.3 | 90.3 | 85 | No |
| Bone diseases adjusted | 0.888 | 99.3 | 89 | 84.3 | No |
| Thyroid conditions adjusted | 0.891 | 99.3 | 89 | 85 | No |
| Heart conditions adjusted | 0.898 | 99.3 | 90.7 | 85 | No |

**Table S2c: *MCC and correctly classified samples for unadjusted and confounder adjusted models comparing PD participant at recruitment and at one year follow up.***

| Model | Overall MCC | Correct PD Year 1 classification (%) | Correct PD Year 2 classification (%) | Confounder in top 50% variables? |
| --- | --- | --- | --- | --- |
| Original | 0.833 | 92.5 | 90.8 | n/a |
| Levodopa LED adjusted | 0.833 | 92.5 | 90.8 | No |
| LLED (mg) adjusted | 0.833 | 92.5 | 90.8 | No |
| MDS_UPDRS Score adjusted | 0.833 | 92.5 | 90.8 | No |
| MOCA Score adjusted | 0.833 | 92.5 | 90.8 | No |
| HY stage adjusted | 0.838 | 92.5 | 91.2 | No |

**Table S3: *For classification and regression models when confounder used as outcome, Matthew Correlation Coefficient (MCC) and R2, used as performance indicators.***

|  | PD vs Control | PD v iRBD v Control |
| --- | --- | --- |
| Gender MCC | 0.345 | -0.063 |
| High cholesterol MCC | 0.067 | -0.009 |
| Hypertension MCC | 0.015 | 0.002 |
| Skin condition MCC | -0.011 | n/a* |
| Bone disease MCC | -0.028 | -0.051 |
| Thyroid condition MCC | 0 | -0.039 |
| Heart condition MCC | 0 | -0.021 |
| Age R2 | -0.091 | -0.222 |
| BMI R2 | -0.107 | -0.214 |
| Alcohol R2 | -0.244 | -0.193 |
| Smoking R2 | -0.154 | -0.125 |

**indicates not sufficient samples per group to perform analysis.*

**Table S4: Prediction of clinical characteristic and drug dosage/intake within PD cohort (from Manchestger cohort the data taken is from visit one only) using measured volatilome. MCC and R^2^ used for model performance measurement.**

| Clinical characteristic or medication | Longitudinal PD comparison | PD only UK | PD only UK and Austria |
| --- | --- | --- | --- |
| HY Stage MCC | 0.137 | -0.04 | n/a |
| MOCA Score R^2^ | -0.174 | -0.099 | n/a |
| MDS_UPDRS Score R^2^ | -0.307 | -0.261 | n/a |
| LLED (mg) R^2^ | -0.455 | -0.304 | n/a |
| Levodopa dosage R^2^ | -0.493 | -0.286 | n/a |
| Amantadine MCC | n/a | 0 | -0.019 |
| Pramipexole MCC | n/a | -0.045 | -0.009 |
| Ropinirole MCC | n/a | -0.017 | -0.037 |
| Rotigotone MCC | n/a | 0 | 0 |
| Rasagiline MCC | n/a | 0.284 | 0.182 |
| Entacapone MCC | n/a | n/a* | -0.023 |
| L-dopa MCC | n/a | -0.113 | 0.328 |
| Comorbidity MCC | n/a | 0.143 | n/a |
| NDS R^2^ | n/a | -0.093 | n/a |
| VAS R^2^ | n/a | -0.228 | n/a |
| mins since last medication R^2^ | n/a | -0.541 | n/a |
| Tremor Score R^2^ | n/a | -0.16 | n/a |
| PIGD Score R^2^ | n/a | -0.314 | n/a |
| Tremor/PIGD ratio R^2^ | n/a | -0.258 | n/a |
| Years of PD symptoms R^2^ | n/a | -0.442 | n/a |
| Years since diagnosis R^2^ | n/a | -0.398 | n/a |
| Months since diagnosis R^2^ | n/a | -0.398 | n/a |
| Tremor or PIGD or Intermediate MCC | n/a | -0.211 | n/a |

*n/a indicates no data available for the cohort and * indicates not sufficient data for samples in each class for analysis.*
